# Molecular characteristics of phages located in Carbapenemase-Producing *Escherichia coli* clinical isolates: New Phage-Like Plasmids

**DOI:** 10.1101/2022.06.29.498070

**Authors:** Maria Lopez-Diaz, Ines Bleriot, Olga Pacios, Laura Fernandez-Garcia, Lucia Blasco, Anton Ambroa, Concha Ortiz-Cartagena, Neil Woodford, Matthew J. Ellington, Maria Tomas

## Abstract

*Escherichia coli* normally inhabits the gastrointestinal tract of humans and animals. Most *E. coli* bacteria do not cause problems, but the acquisition of different resistance and virulence genes encoded by mobile plasmids or phages by different bacterial isolates has been associated with the appearance of successful high-risk clones of multidrug-resistant (MDR) *E. coli* such as ST131 or ST405. In the present study, 50 temperate bacteriophages present in 21 clinical isolates of carbapenemase-producing *E. coli* of sequence types (STs) ST38, ST131, ST167, ST405 and ST410 were analysed. These phages were classified in the three families of the order *Caudovirales*: 24 within the family *Siphoviridae*, 23 in *Myoviridae* and 3 in *Podoviridae*. The size of the phages studied ranged from 11 to 95 Kb. Phylogenetic analysis of the terminase large subunit allowed us to classify these phages into different groups showing similarity with the phage sequences deposited in the Microbe Versus Phage (MVP) database and which belonged to clusters 229, 604, 2503 and 2725. On the other hand, bioinformatic study revealed that most of the identified proteins exerted a structural function (26.73%) but also functions involved in lysis/lysogeny (6.70%) or regulation (5.20%) among others. In addition, the ParA-ParB partitioning system and the type II toxin-antitoxin Phd-Doc system were also found in two of the phages studied, which could indicate the presence of plasmid-prophages. Host range testing revealed that two isolates were more susceptible to infection than the other isolates.

**IMPORTANCE:** *Escherichia coli* is one of the pathogens that causes most problems in human health, as it presents multiple resistances to different antibiotics. The study of bacteriophages located in different isolates of this species is important for the development of new anti-infective therapies. Currently, antibiotic resistance is a major problem, but more and more studies are pointing to experimental treatments with bacteriophages as a possible solution.

## INTRODUCTION

*Escherichia coli* has emerged as one of the pathogens with the most epidemiological implications, causing a multitude of infections, being the most frequent agent of urinary tract infections and the main cause of bacteremia due to gram-negative bacteria (1). In addition to being part of the intestinal microbiota, *E. coli* can survive in both sewage and drinking water, plants, animals and food (2). It is capable of easily capturing genetic material by horizontal gene transfer (HGT) such as transposons, plasmids or phages encoding different virulence factors. Most *E. coli* bacteria are commensal and therefore harmless, but the acquisition of different resistance mechanisms has been associated with the emergence of high-risk clones that disseminate readily, such as ST131 or ST405 (3, 4).

In addition, antibiotic usage has also contributed to an increase in the incidence of antibiotic resistance. This has led to the need to develop new strategies as therapeutic alternatives, one of which is the use of bacteriophages (phages), commonly known as “phage therapy” (5).Discovered more than a century ago, phages exclusively infect bacteria and are the most abundant biological entities on the planet, and are also found in our own microbiota. Most phages have a high host specificity, which allows them to recognize and lyse specific bacterial isolates, thus rendering them harmless to humans. Others are capable of recognizing a wide range of closely related hosts within the same species. In this regard, phage therapy is proposed as an useful tool (6).

In order to study the evolutionary relationships and phylogeny of prophages, we analyzed the genomic characteristics of 50 intact prophages detected in 21 clinical isolates of carbapanemase-producing *E. coli* (CP *E. coli*).

## RESULTS

### Prophage prediction and classification

In the present study, the genome sequences of 21 CP *E. coli* clinical isolates were analysed for prophage identification using PHASTER bioinformatics tool (http://phaster.ca/) (Table 2). A total of 165 prophage-like elements were detected of which 50 were classified as intact, 20 as questionable and 95 as incomplete. The highest number of prophages, six, occurred in Eco_569 strain and none intact prophage region was found in ST410 (Figure 1A). The fact that no prophage was found does not mean that they are not present, but the PHASTER software could be limiting in this detection as working with contigs makes it difficult to detect complete phages, in fact incomplete or questionable prophages in ST410 were found but they were not included in the present study. Families were assigned by sequence homologies with the most common bacteriophages listed by PHASTER in the Virus-Host Database. (https://www.genome.jp/virushostdb/). All 50 prophages belonged to the order *Caudovirales*. Most were members of the family *Siphoviridae*: 24, and the others belonged to the families *Myoviridae*:23 and *Podoviridae*:3 (Figure 1B).

**Figure 1.**
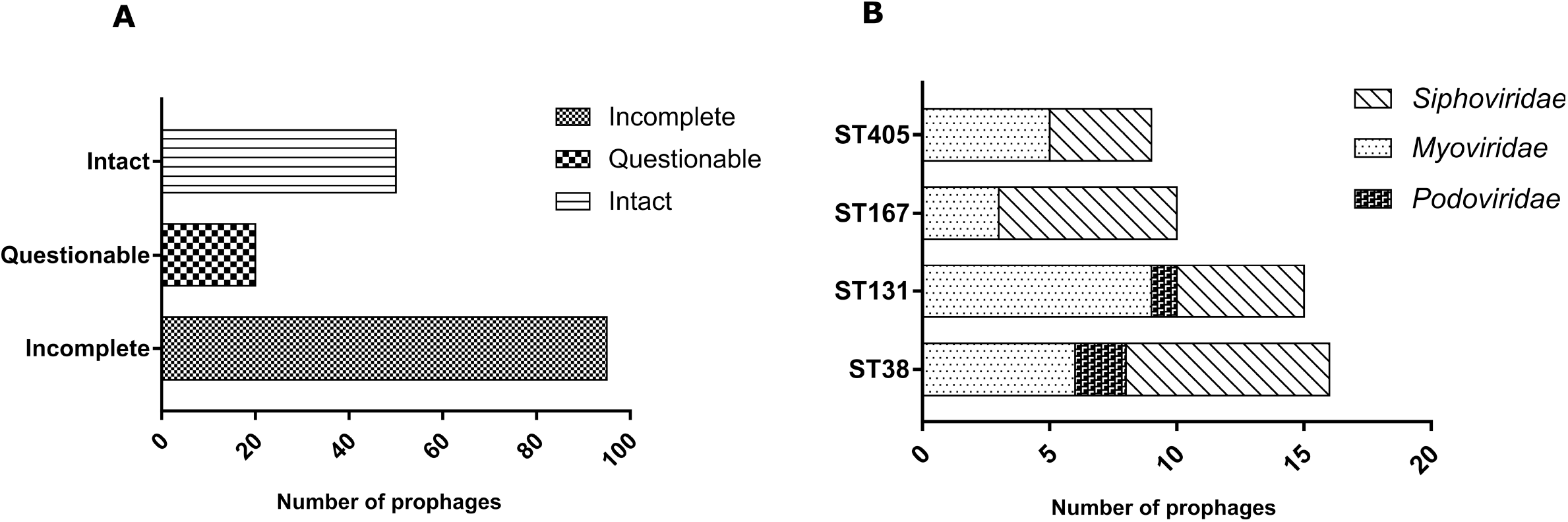
(A) Bars graph showing the distribution of total prophages (intact, incomplete, and questionable) in *E. coli* isolates. (B) Distribution of intact prophages in *E*.*coli* genomes by family.

**Figure 2.**
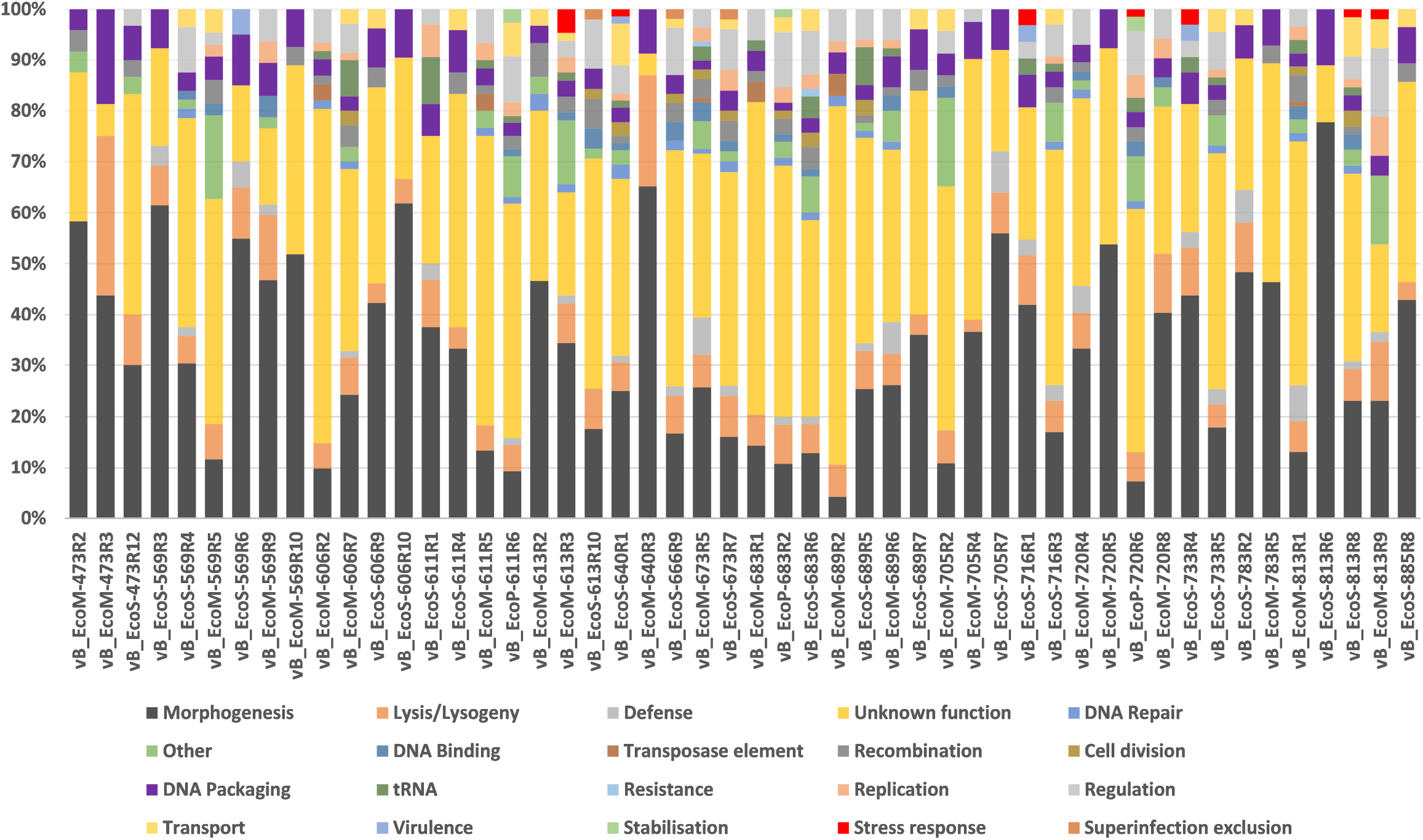
Functional classification of proteins encoded by prophage regions in CP *E. coli* genomes.

**Figure 3.**
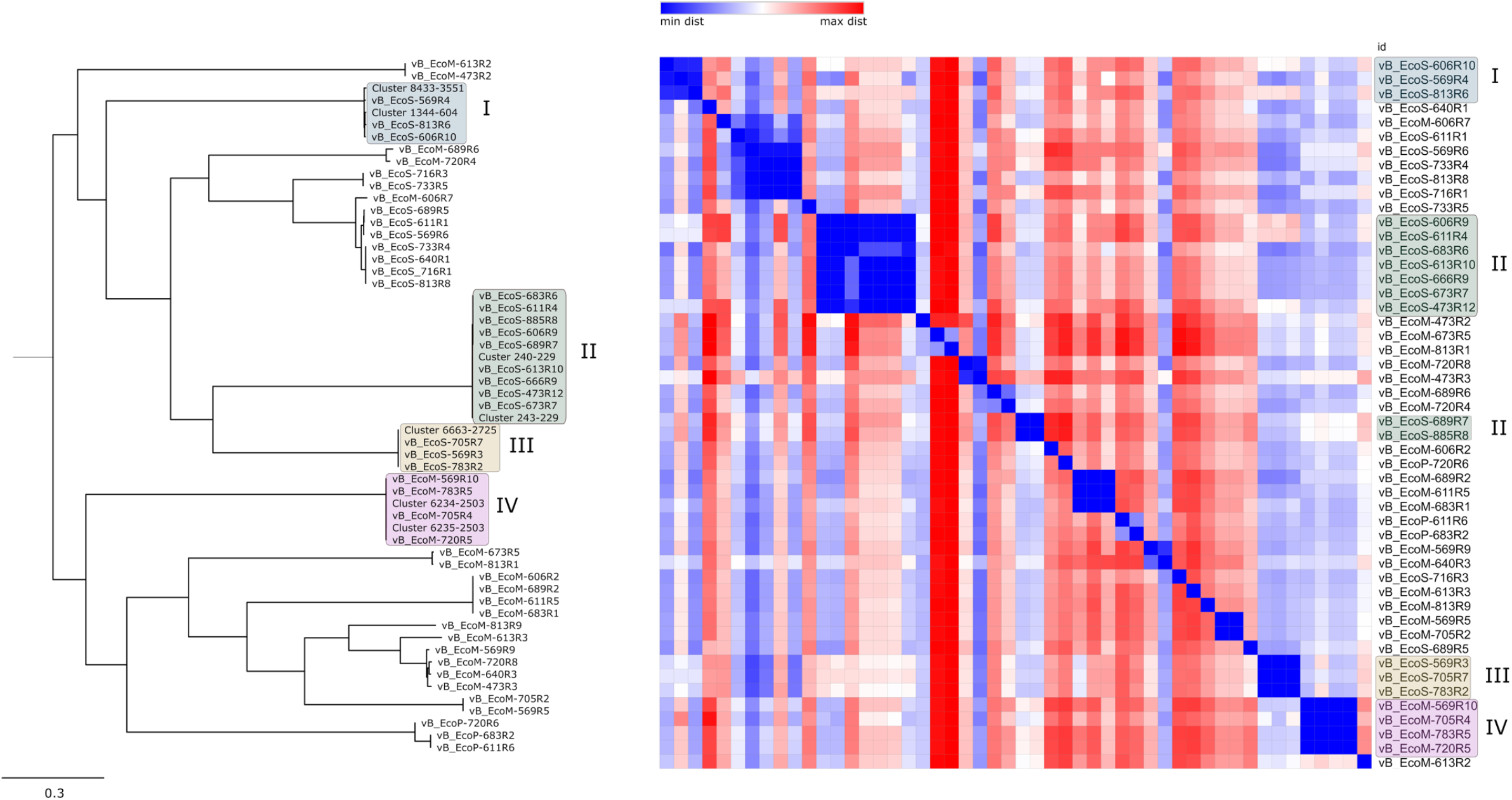
Phylogeny of the large terminase subunit and SNP analysis. Pairwise SNP distance matrix from a FASTA sequence alignment.

### Prophage Annotation

The most abundant functional category found in prophage genomes are genes related to morphology (26.73%), lysis/lysogeny (6.70%), regulation (5.20%) or DNA packaging (4.39%) and recombination (3.30%). One of the proteins identified was terminase which enables viral DNA packaging. The large subunit of the terminase was used to analyze the phylogenetic relationships between the phages in this study since it was present in each of them, unlike the rest of the proteins or even the small subunit that could only be found in some prophages.

Proteins involved in host cell lysis were identified, such as endolysins which were found in 23 of the 50 phages analyzed or holins were found in 36 phages. Additional proteins called spanins were also found in 28 phages. Those three proteins were usually found together in the same phages. On the other hand, proteins involved in the integration of viral DNA into the host, i.e. those involved in lysogeny processes, such as integrase, were found in only 20 phages. In addition, a large number of HNH homing endonucleases were identified.

Regarding the defense systems present in prophages, we can highlight the presence of the type II toxin-antitoxin system PhD-Doc in the phages vB_EcoM-673R5 and vB_EcoM-813R1. This is one of the smallest families of this type of systems and was first discovered in the bacteriophage P1 of *E. coli* (7).

The endodeoxyribonuclease toxin RalR was also found in the prophages vB_EcoS-666R9, vB_EcoS-673R7 and vB_EcoS-683R6. This is a toxic component of a type I toxin-antitoxin (TA) system.

The presence of the transcription regulator genes cI and Cro was observed in most of the prophages, which play a key role in their life cycle.

tRNA-scan-SE identified tRNAs only in 14 of the 50 genomes (Table 2) and the phage-bacteria junctions, attL (left) and attR (right) were identified by PHASTER only in 18 of the 50 genomes: vB_EcoM-689R6, vB_EcoP-611R6, vB_EcoP-683R2, vB_EcoS-683R6, vB_EcoM-569R5, vB_EcoM-569R9, vB_EcoP-720R6, vB_EcoM-720R8, vB_EcoM-705R2, vB_EcoS-733R5, vB_EcoS-640R1, vB_EcoS-813R8, vB_EcoM-813R9, vB_EcoS-716R3, vB_EcoM-613R3, vB_EcoM-613R10, vB_EcoS-666R9, vB_EcoS-673R7. In the rest of phage sequences no attL/R sites or termini regions were identifiable, this may be due to contig boundaries that may negatively affect predicted prophage scores, as it results in genes or regions that are ignored because they are not in the contig.

Plasmid partitioning systems ParA-ParB associated to site-specific tyrosine recombinase Cre were identified in two phages (vB_EcoM-673R5 and vB_EcoM-813R1). This may indicate a plasmid prophage stage (8).The recombinase Cre is derived from the *Enterobacteria*phage P1 and its role is to maintain the phage genomes as a monomeric unit-copy plasmid in the lysogenic stage. In addition, the ParB partition protein without ParA or the recombinasae Cre was localized in other phages (vB_EcoM-473R2, vB_EcoS-473R12, vB_EcoM-569R10, vB_EcoS-606R9, vB_EcoS-611R4, vB_EcoM-613R2, vB_EcoS-613R10, vB_EcoS-666R9, vB_EcoS-673R7, vB_EcoS-683R6, vB_EcoS-689R7, vB_EcoS-716R3, vB_EcoM-720R5, vB_EcoS-733R5, vB_EcoM-783R5, vB_EcoS-885R8) but in this case does not necessarily indicate that the phage has a plasmid prophage stage.

Besides, the virulence-associated gene *bor* was identified in phages vB_EcoS-569R6, vB_EcoS-640R1, vB_EcoS-716R1, vB_EcoS-733R4 and vB_EcoS-813R8. *bor* is a phage gene that are expressed during lysogeny and whose products show homology with bacterial virulence proteins (9). CRISPR-related or anti-CRISPR proteins were not found in the present work.

### Prophage Phylogenetic Analysis

Phylogenetic analysis of the long terminase subunit of the 50 intact phages together with representative MVP clusters was performed. In addition, a heatmap of phylogenetic distances (number of single nucleotide polymorphisms (SNPs) was also performed using the complete phage sequences. A correlation was observed between the clusters formed with the long terminase subunit and the MVP clusters and the clusters formed by SNP analysis.

### Comparative genomic analysis of *E. coli* temperate bacteriophages

Once the annotation of the prophages with RAST and HMMER (phmmer) was performed, the existence of phages of identical or similar size was observed in different isolates. In isolates Eco_716 and Eco_733, the same phage with a size of 46651 bp was found. In other isolates, phages of very similar size and distribution were found. The comparative analysis of these phages was performed using the BRIG program, in which the complete sequences of the phages as well as the sequences of prophages obtained from the MVP database were used.

Amongst one grouping of phages, 5 phages showed a high similarity with the sequence of the reference phage 1344 of cluster 604 of the MVP database (Figure 4). In a second group (II), two subgroups were observed, in subgroup II-A (Figure 5) five phages were most similar to sequence 240 of cluster 229, one phage (EcoS_473R12) was larger in size but shared most of its genome with the other phages in the group. In subgroup II-B (Figure 6), phages showed similarity with sequence 243 of cluster 229. In this case, the sequence of the cluster was smaller but the region that was present was similar to the 3 phages included. In group III (Figure 7), sequence 6663 of cluster 2725 was very similar to the phage EcoS_783-R2 and, albeit to a lesser extent, Eco_569R3. In group IV (Figure 8), we observed that the sequence 6234 of cluster 2503 presented high similarity to one phage EcoM_705R4, both in size and genomic content, whilst the rest of the phages in this group presented a smaller size.

**Figure 4.**
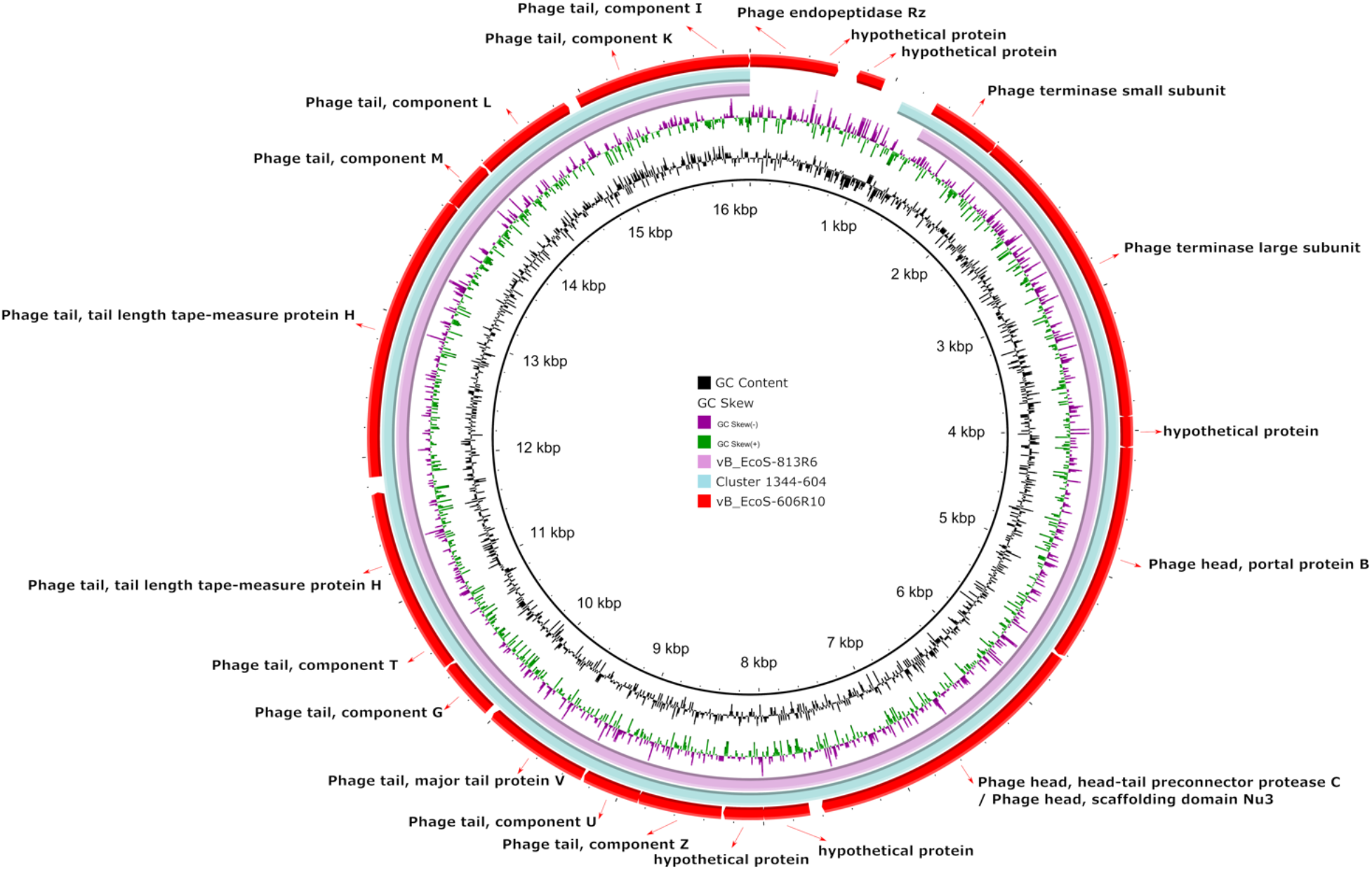
Comparative genomic analysis of CP *E. coli* temperate bacteriophages with cluster 1344-604 (query cover >99.9 %; identity >99.8 %) from the MVP database (https://mvp.medgenius.info) was performed using the BRIG software.

**Figure 5.**
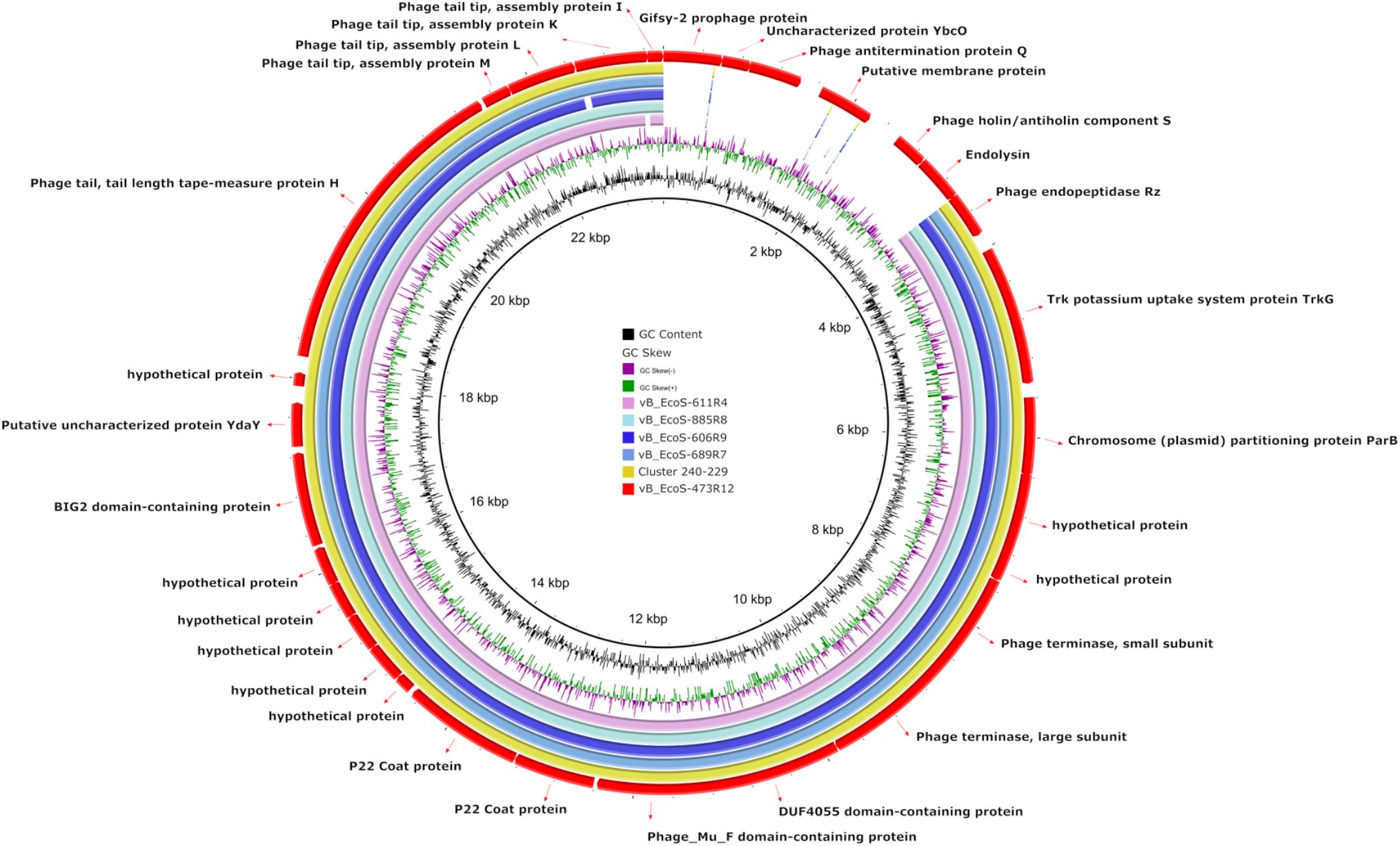
Comparative genomic analysis of CP *E. coli* temperate bacteriophages with cluster 240-229 (query cover >99.9 %; identity >99.8 %) from the MVP database was performed using the BRIG software.

**Figure 6.**
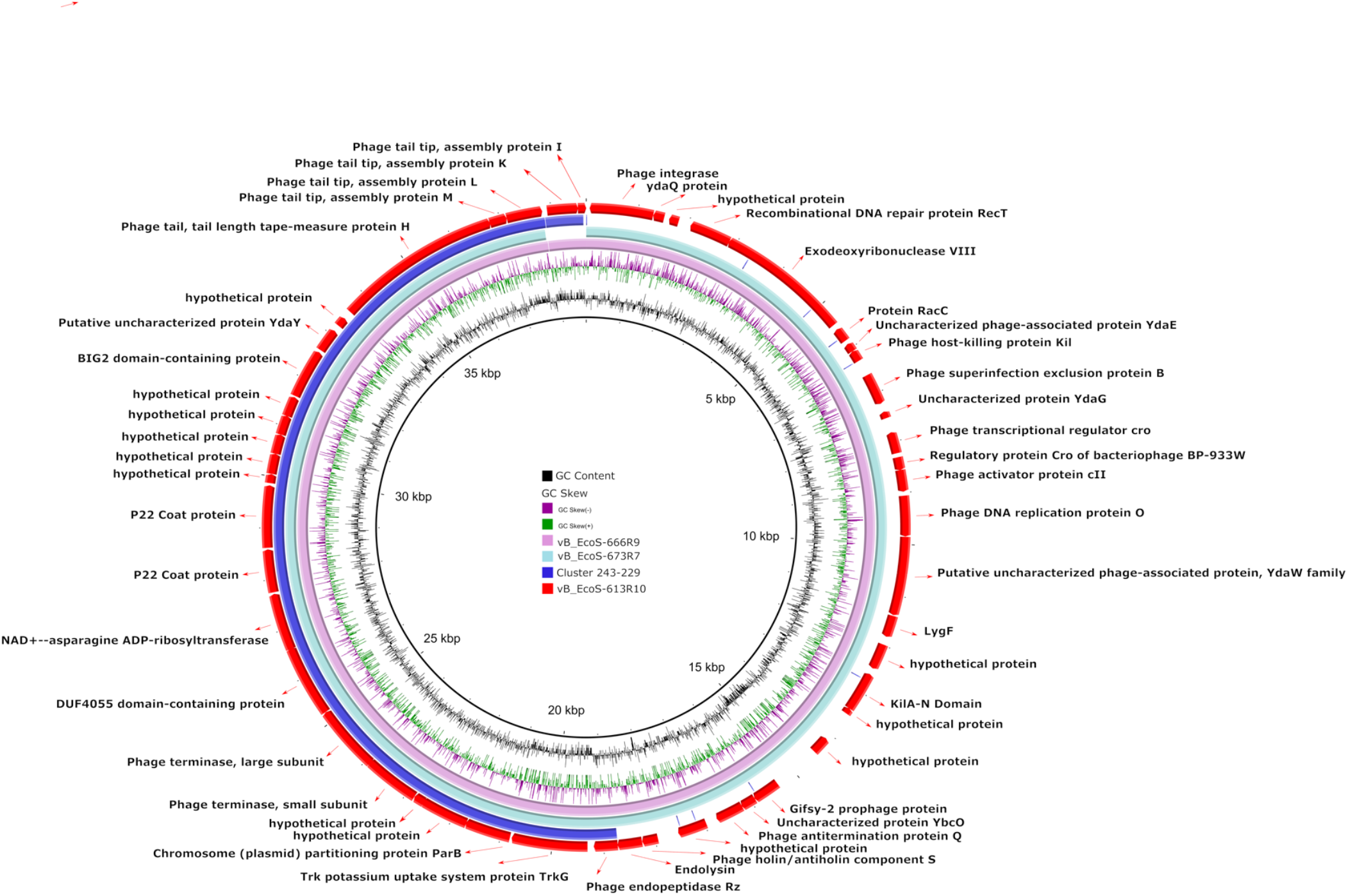
Comparative genomic analysis of CP *E. coli* temperate bacteriophages with cluster 243-229 (query cover >99.9 %; identity >99.8 %) from the MVP database was performed using the BRIG software.

**Figure 7.**
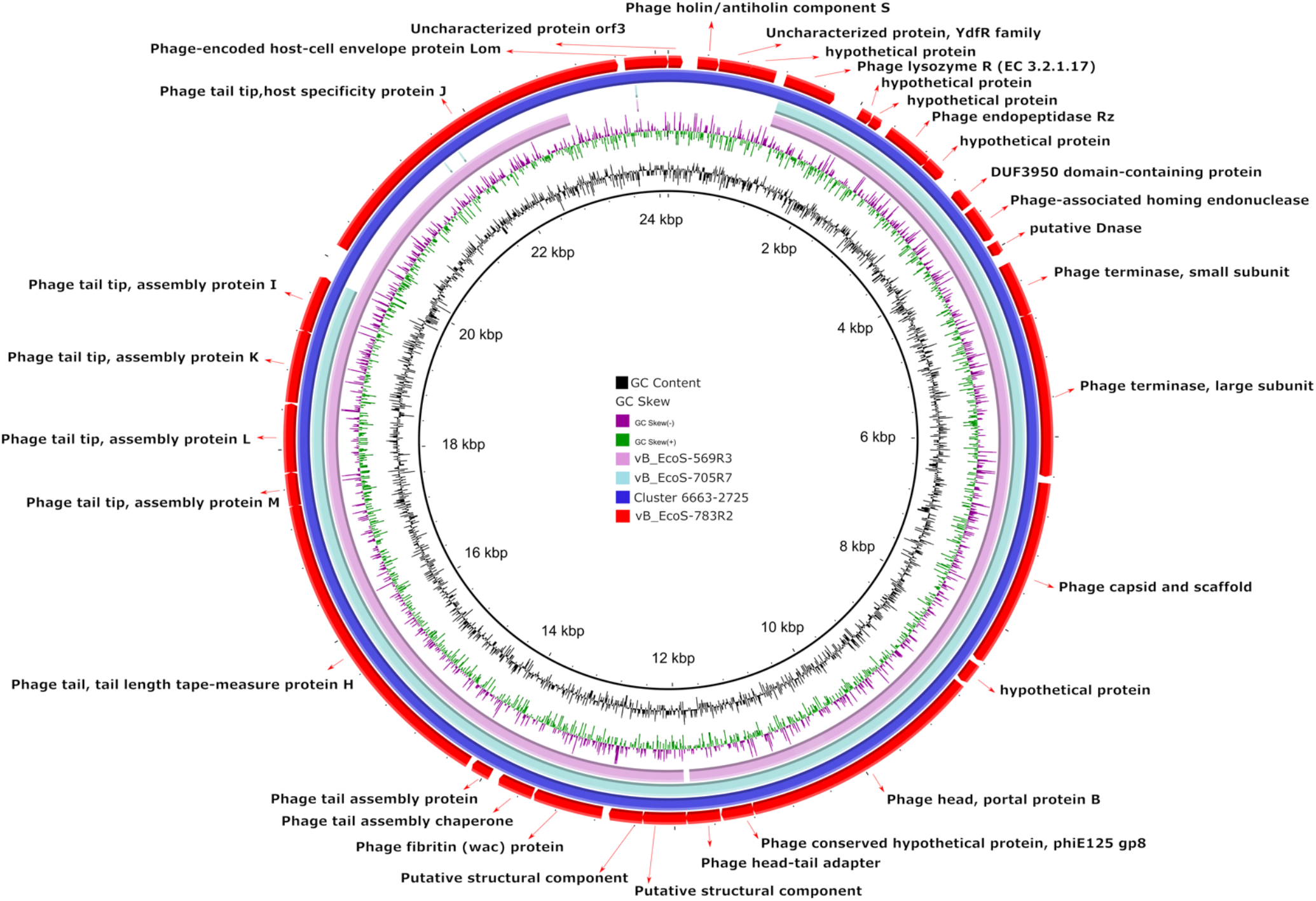
Comparative genomic analysis of CP *E. coli* temperate bacteriophages with cluster 6663-2725 (query cover >99.9 %; identity >99.8 %) from the MVP database was performed using the BRIG software.

**Figure 8.**
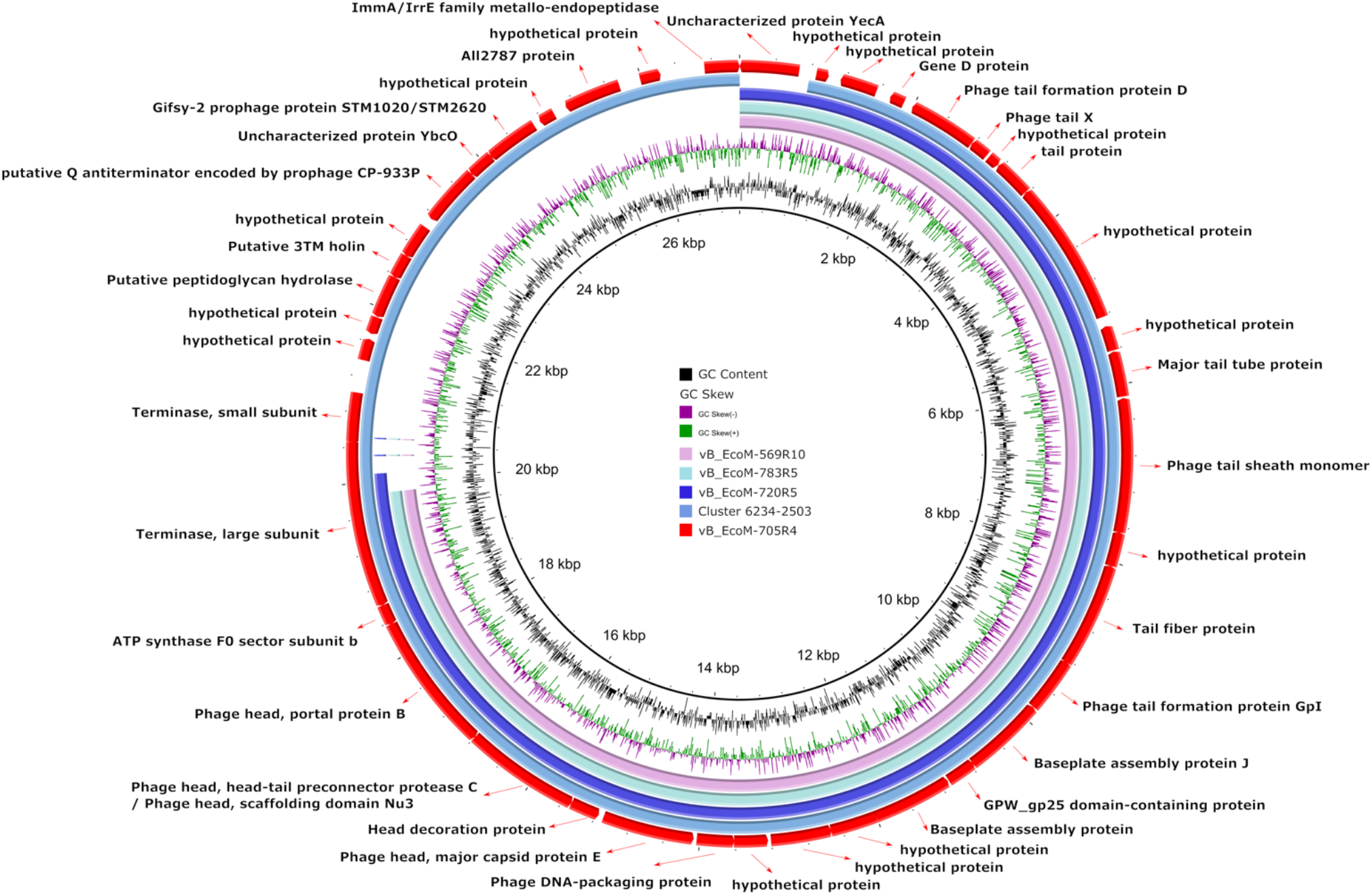
Comparative genomic analysis of CP *E. coli* temperate bacteriophages with cluster 6234-2503 (query cover >99.9 %; identity >99.8 %) from the MVP database was performed using the BRIG software.

### TEM studies

The three families belonging to the order *Caudovirales, Siphoviridae, Myoviridae* and *Podoviridae*, were observed in the TEM images (see Figure 9 for representatives).

**Figure 9.**
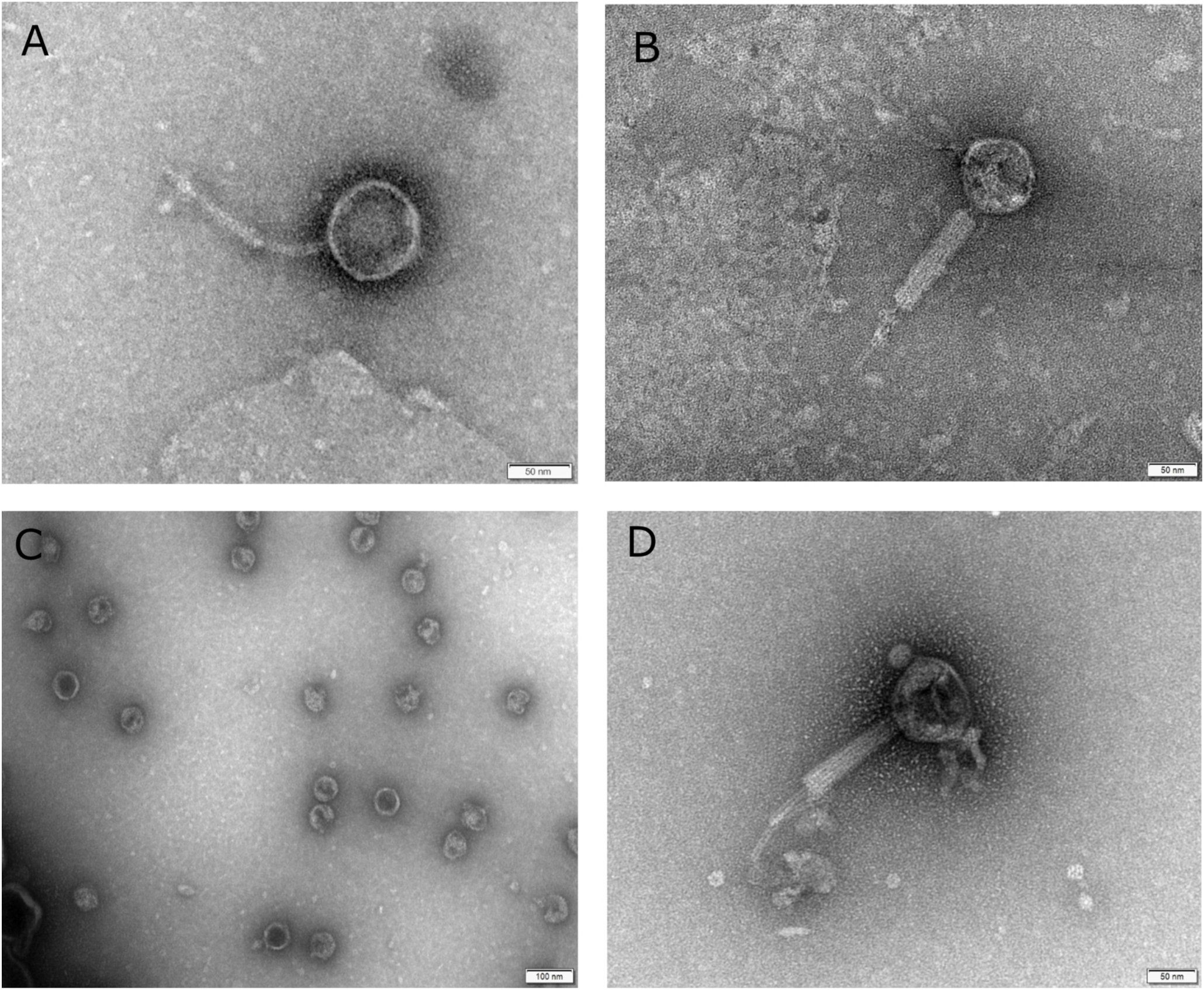
TEM images showing the different families of prophages present in different clusters. (A)The family *Siphoviridae* obtained from Eco_611 (B and D). The family *Myoviridae* from Eco_473 and Eco_569 respectively (C). The family *Podoviridae* from Eco_60.

### Plating Analyses for Determination of Host Range

Host range assay of eight mixtures of induced bacteriophages isolated from eight representative clinical *E. coli* isolates (Eco_683 (1), Eco_720 (2), Eco_569 (3), Eco_611 (4), Eco_813 (5), Eco_716 (6), Eco_606 (7) and Eco_473 (8)) and present in each of the 4 identified clusters. The infectivity of these cocktails was tested in the 21 isolates of CP *E. coli* studied. The isolates infected by each phage mix are shown in green and those that are not infected in red. Interestingly, two of the isolates, Eco_689 and Eco_613, were found to be more susceptible to infection by a wider range of mixtures of bacteriophages.

### Data summary

The genome sequences of the clinical isolates of *Escherichia coli* and temperate bacteriophages have been deposited in GenBank under BioProject accession number PRJNA834992 (http://www.ncbi.nlm.nih.gov/bioproject/834992).

## DISCUSSION

In the present study, 50 bacteriophages from 21 clinical isolates of CP *E. coli* were analysed and revealed a wide genomic diversity of prophages. It was considered necessary to carry out an in-depth analysis of the phage genome in order to study their phenotypic characteristics, their evolution and distribution. To date, most studies were focused to identify prophages on the enviroment, such as in wastewater (10) or on Shiga toxins (Stx) of Shiga toxin-producing *E. coli* (STEC) which are generally encoded in the genome of lambdoid bacteriophages causing of numerous food-borne outbreaks and serious diseases such as haemorrhagic colitis (11, 12).

To our knowledge, this is the first study that aims to analyse different phages present in carbapenemase-producing *E. coli* clinical isolates that also belong to high-risk clones. We used a high number of genome sequences to provide a more detailed and comprehensive prophage distribution.

The order *Caudovirales* is considered to be probably older than the separation of life into the three-domain system (Bacteria, Archaea and Eukarya). This has led to various interactions between viruses and bacteria, such as lysogeny in which viruses are able to integrate into the bacterial genome and may increase both host fitness and prophage survival; or lysis leading to bacterial death (13). Through lysogeny, phages and bacteria can be protected in the environment and they are able to exchange material between hosts such as resistance or virulence genes (14).In the present study, the *bor* gene was detected in several phages. *bor* is closely related to the iss locus of the ColV,I-K94 plasmid (9). This gene is closely related to the iss locus of the ColV,I-K94 plasmid, which promotes bacterial resistance to serum complement elimination in vitro and virulence in animals. According to previous studies, it was also found in other lamboid coliphages and in several Extraintestinal pathogenic *E. coli* (ExPEC) clinical isolates. Therefore, due to its high prevalence, virulence and the fact that it is transmitted in mobile elements, an in-depth study of both this gene and the protein it produces would be necessary as they could be useful diagnostic tools or vaccine targets for the prevention of various diseases caused by ExPEC (15).

On the other hand, the presence of the ParA-ParB partitioning system and its association with Cre recombinase in the two phages vB_EcoM-673R5 and vB_EcoM-813R1 could indicate the presence of a plasmid prophage. However, the presence of these proteins alone does not guarantee that the phage replicates as a plasmid (8). Cre recombinase is derived from the *Myoviridae* phage P1 of *E. coli* K12 (the first circular plasmid prophage to be discovered (16)) and its function is to maintain the phage genomes as a single-copy monomeric plasmid in the lysogenic stage. BLASTn alignment of the phages displaying the above-mentioned partitioning system, which could function as a plasmid phage, with *E. coli* K12 phage P1 was performed and showed 76% coverage and 98.68% identity, respectively, for phage vB_EcoM-673R5; and 88% coverage and 96.92% identity, respectively, for phage vB_EcoM-813R1, which could indicate that these phages could act as plasmid prophage but further analysis would be necessary to verify this function.

In addition, the type II TAs PhD-Doc was also identified in these same phages. This is one of the smallest families of such systems and was also first discovered in the same bacteriophage mentioned above, bacteriophage P1 of *E. coli* (17).This class of TAs is widely represented in mobile genetic elements such as plasmids and phages, but also in bacterial chromosomes (18). In our case, this could further strengthen the theory that these two phages detected in the present study would function as plasmid prophages.

Other proteins identified were a large number of HNH homing endonucleases, which play a key role in DNA packaging and thus in the phage life cycle as previous studies described (19, 20). The endodeoxyribonuclease toxin RalR was also found in the prophages vB_EcoS-666R9, vB_EcoS-673R7 and vB_EcoS-683R6. This is a toxic component of a type I toxin-antitoxin (TA) system. When overexpressed, it inhibits bacterial growth and the cells become filamentous. The toxic effects would be neutralized by RalA antitoxin, but in the present study it was not detected (21).

The presence of the transcription regulator genes cI and Cro was observed in most of the prophages, which play a key role in their life cycle. When cI proteins predominate, the phage remains in a lysogenic state, whereas when the Cro protein increases, it will go into a lytic state (22). When this phase is activated, disruption of each of the three layers (inner membrane, peptidoglycan and outer membrane) of the cell envelope occurs. In addition to endolysins and holins, in the case of Gram-negative bacteria, as *E. coli*, spanins are necessary to disrupt the outer membrane (23, 24). In the present study, the endolysin-holin-spanin complex, necessary for effective lysis, was detected in more than half of the phages.

In recent years, endolysins (also known as enzibiotics) have emerged as a promising alternative to antibiotics in the treatment of infections caused by multi-resistant bacteria. A recent study has demonstrated the broad spectrum of action of the endolysin ElyA1 that would allow more MDR Gram-negative bacteria to be included as targets for antimicrobial treatment combined with colistin (25).

In order to study the distribution of the phages, the phage sequences were compared, using BLASTn, with the sequences deposited in the MVP database. Some of the phages showed an identity and coverage of more than 95% with some of the sequences grouped in the MVP clusters. This allowed us to classify our phages into different groups (I, II, II and IV). The group that grouped the most phages was group II, which included cluster 229, one of the clusters that grouped the most sequences in the MVP database with 71 *E. coli* sequences.

Interestingly, host range analysis showed that two isolates (Eco_689 and Eco_613) were more likely to be infected than the rest of the study isolates. These results suggest that the number of *E. coli* isolates capable of supporting prophages development is low. This could mean that resistant bacteria must possess another type of mechanism, unrelated to prophages, that allows them to resist infection by phages. This highlights the need for in-depth studies of not only the phage carrying those isolates but the entire genome of the bacterium.

## MATERIAL AND METHODS

### *Escherichia coli* isolates

In the present study, we analysed 21 CP *E. coli* clinical isolates submitted to Public Health England’s (PHE) Antimicrobial Resistance and Health Care Associated Infections (AMRHAI) Reference Unit. Those isolates were collected from different English Regions: London, East of England, South East, West Midlands, North West and Scotland and belong to 5 different sequence types (STs): 38, 131, 167, 405 and 410, considered “high-risk” clones (Table 1).

**Table 1.** Characteristics of the *E. coli* clinical isolates.

| Isolate | MLST | Isolation source | Collection year | Region | GenBank accession number |
| --- | --- | --- | --- | --- | --- |
| Eco_606 | 38 | Urine | 2015 | Scotland | SAMN28088462 |
| Eco_689 | 38 | Urine | 2015 | East | SAMN28088463 |
| Eco_611 | 38 | Swab | 2015 | London | SAMN28088464 |
| Eco_683 | 38 | Urine | 2015 | North West | SAMN28088465 |
| Eco_885 | 38 | Urine | 2015 | South East | SAMN28088466 |
| Eco_569 | 131 | Screening | 2015 | London | SAMN28088467 |
| Eco_720 | 131 | Sputum | 2015 | West Midlands | SAMN28088468 |
| Eco_705 | 131 | Screening | 2015 | London | SAMN28088469 |
| Eco_783 | 131 | Urine | 2015 | North West | SAMN28088470 |
| Eco_733 | 167 | Screening | 2015 | London | SAMN28088471 |
| Eco_640 | 167 | Screening | 2015 | East | SAMN28088472 |
| Eco_813 | 167 | Screening | 2015 | West Midlands | SAMN28088473 |
| Eco_716 | 167 | Screening | 2015 | London | SAMN28088474 |
| Eco_673 | 405 | N/A | 2015 | Scotland | SAMN28088475 |
| Eco_473 | 405 | Screening | 2015 | London | SAMN28088476 |
| Eco_613 | 405 | Screening | 2015 | London | SAMN28088477 |
| Eco_666 | 405 | Screening | 2015 | East | SAMN28088478 |
| Eco_684 | 410 | Urine | 2015 | London | SAMN28088479 |
| Eco_766 | 410 | Sputum | 2015 | London | SAMN28088480 |
| Eco_703 | 410 | Urine | 2015 | South East | SAMN28088481 |
| Eco_783_410 | 410 | Urine | 2015 | North West | SAMN28088482 |

**Table 2.**
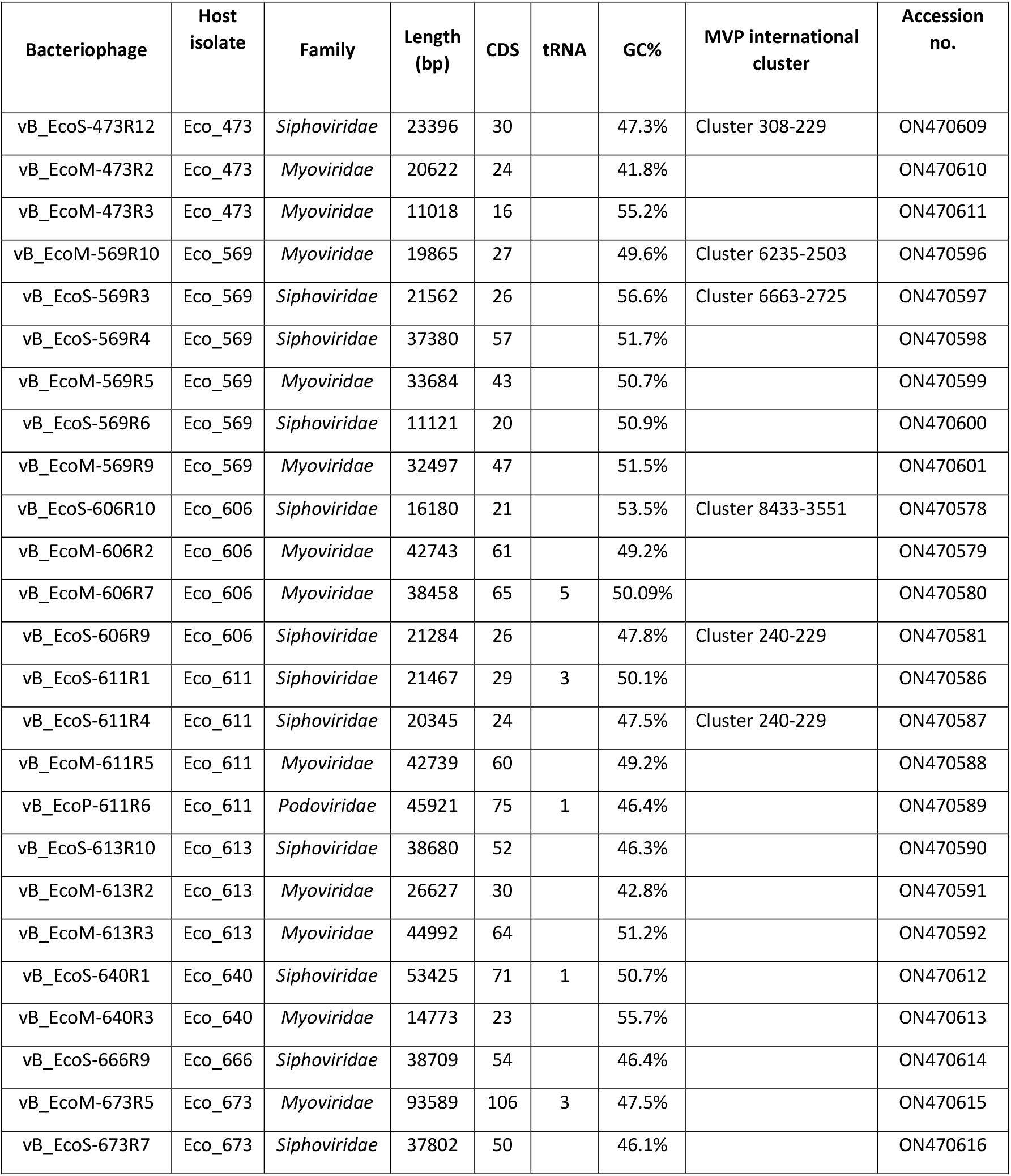
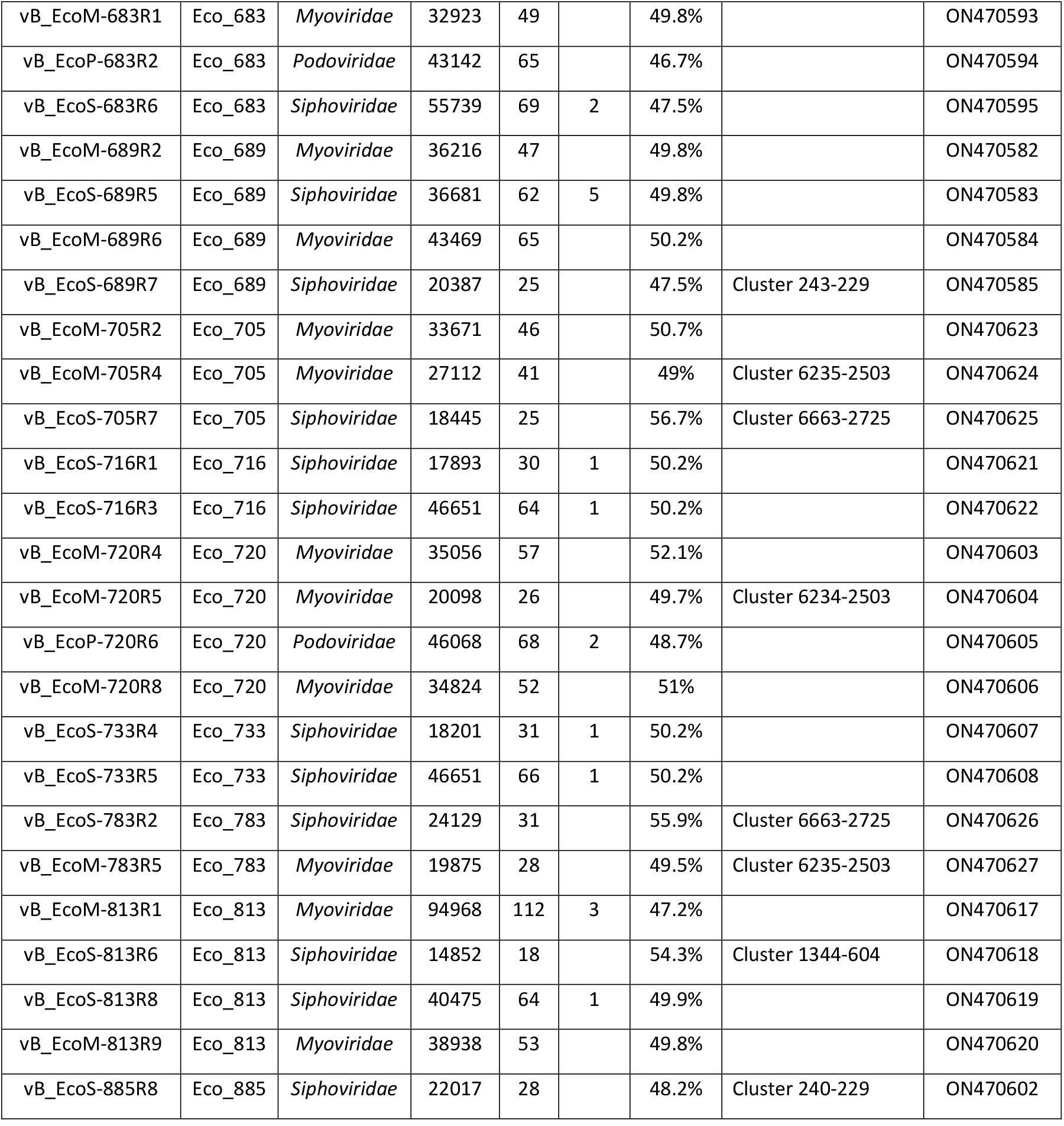
Characteristics of the genome sequences of the 50 prophages found in 21 clinical isolates of CP *E. coli*.(-) Not detected

**Table 3.**
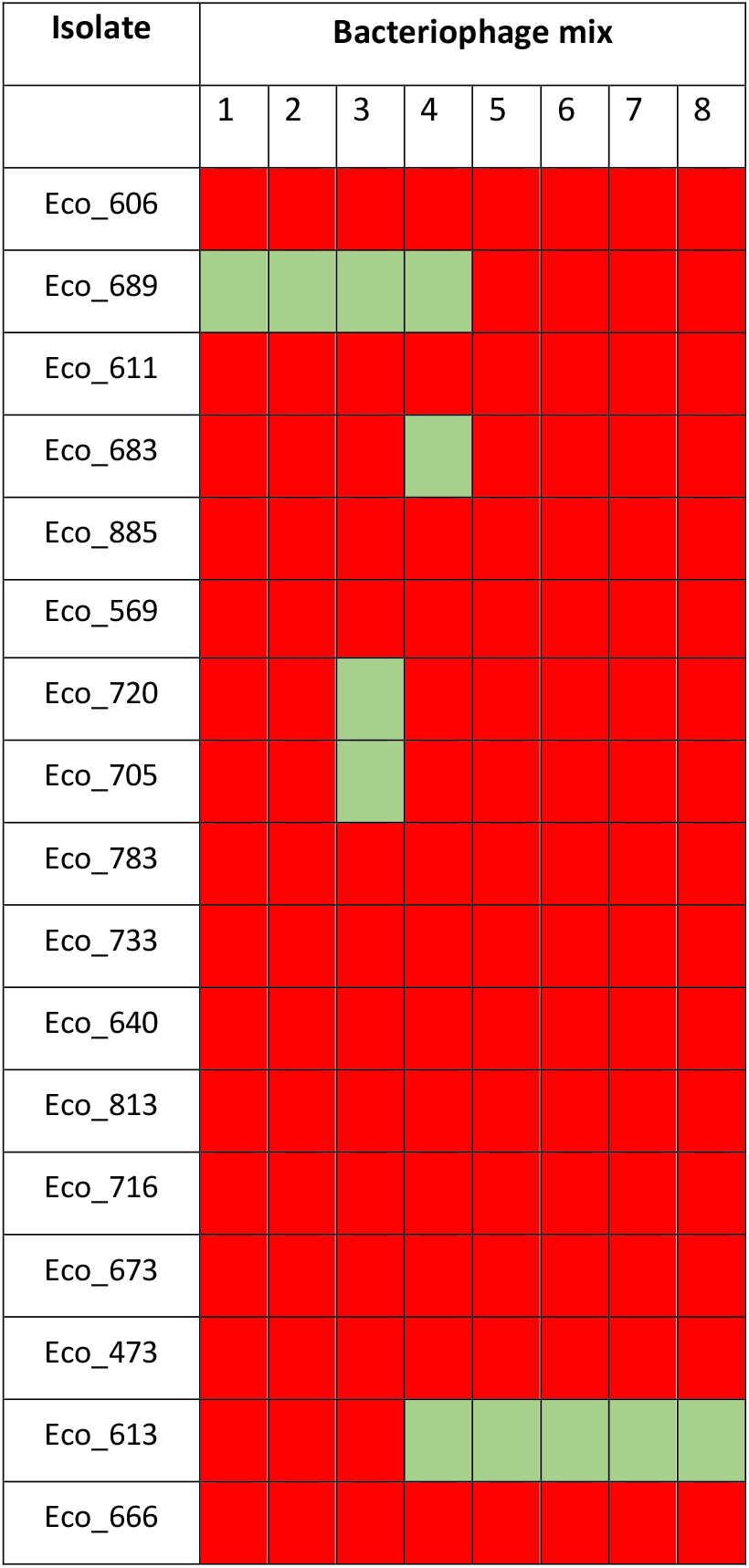
Host range assay

### Genome sequencing

Between 2014 and 2016, AMRHAI performed whole genome sequencing of carbapenemase-producing *E. coli* isolates using the QIAsymphony DSP DNA Midi kit (Qiagen GmbH, Hilden, Germany) and the Nextera XT DNA library prep kit (Illumina). Sequencing was performed on a HiSeq 2500 (Illumina) and assembly was performed using SPAdes 3.10.1.

### Prophage prediction and classification

PHAge Search Tool enhanced release (PHASTER; http://phaster.ca/) (26) was used to identify prophages in the genomes. Only those identified by the program as intact were used in the present study. Families were assigned by sequence homologies with the most common bacteriophages listed by PHASTER in the Virus-Host Database. (https://www.genome.jp/virushostdb/). In order to analyse bacteriophage-host interaction, MVP (Microbe Versus Phage) database (27) was used to compare *E. coli* prophages with phage clusters, the first 95 most abundant *E. coli* clusters were downloaded. Each prophage was compared to MVP reference phage cluster sequences with >95% identity and coverage using BLASTn. To compare the sequences of prophage sequences of similar length and related MVP groups, the BLAST Ring Image Generator (BRIG) (28) program was used.

### Prophage annotation

Open reading frames (ORFs) were predicted and annotated using Rapid Annotation Subsystem Technology, version 2.0 (RAST; http://rast.nmpdr.org) and the BLAST-protein tool PSI-BLAST developed by the NCBI (https://blast.ncbi.nlm.nih.gov/Blast.cgi). In addition, we used the phmmer tool which uses a hidden Markov model (HMM) to predict protein domains (https://www.ebi.ac.uk/Tools/hmmer/search/phmmer) (29). The tRNAS detected by RAST in some prophages were confirmed with RNAscan-SE software (30). The CRISPRCasFinder (https://crisprcas.i2bc.paris-saclay.fr/) and anti-CRISPRdb (http://cefg.uestc.edu.cn/anti-CRISPRdb) programs were used to search for CRISPR and anti-CRISPR proteins, respectively.

### Prophage phylogenetic analysis and SNP analysis

The phylogenetic relationship bewteen the sequences of large terminase subunit and the clusters included in the MVP database was studied. The sequences were aligned using MAFFT v7.407 (31) default options and phylogenetic analysis was performed in RaxmlHPC-PTHREADS-AVX2 version 8.2.12 (32) under the GTRGAMMA model and 100 bootstrap replicates. FigTree (http://tree.bio.ed.ac.uk/software/figtree/) was used to visualize the phylogenetic tree. The SNP distance matrix was calculated from the alignment of the sequences of the 50 intact prophages identified using snp-dist 0.7 (https://github.com/tseemann/snp-dists).

### Transmission electro microscopy (TEM) studies and host range

For phage induction, an overnight culture of each strain was diluted (1:100) in LB and incubated at 37°C with shaking until optical density reached 0.5 at wavelength 600nm (OD600). Mitomycin C (Sigma-Aldrich) was then added at a final concentration of 10µg/ml. Once the culture appeared clear, (after 2-4 hours), the lysates were centrifugate at 3500 rpm for 10 min, and the supernatant was filtered using filtration through 0.22 μm filter (Merck Millipore, Germany). Finally, the samples were stored at 4°C until processed for TEM (33). The agar overlay method was used to determine the host ranges of the induced mixed bacteriophage samples obtained from mitomycin C induction. Eight isolates were selected for induction, representative members of each group (I, II, III and IV). When the OD600 of each strain to be infected reached 0.6 nm, 200 µl of each of the 17 *E*.*coli* isolates were mixed with 4 ml of soft agar (0.5 % NaCl, 1 % tryptone and 0.4 % agar, supplemented with 10mM CaCl2 and 10mM MgSO4) and 100 µl of each of the mixed phage samples. This mixture was poured onto TA agar plates (0.5 % NaCl, 1 % tryptone and 1.5 % agar, supplemented with 10mM CaCl2 and 10mM MgSO4). Once the soft medium was solidified, the mixture was added to TA agar plates (0.5 % NaCl, 1 % tryptone and 1.5 % agar). A negative control without phage was included.

## ACKNOWLEDGEMENTS

M.L.-D. was funded by a postdoctoral fellowship from the Xunta de Galicia-GAIN (IN606B-2018/008). This study was supported by the UK Health Security Agency (UKHSA; formerly PHE) and via funding provided to the authors via the European Joint Programme on One Health (ARDIG) under grant number 773830 and the Joint Programming Initiative on Antimicrobial Resistance (JPIAMR) and the Medical Research Council (MRC) as part of the ST131 transmission consortium under grant code MR/R002843/1. The authors would also like to thank the Supercomputing Center of Galicia (CESGA) for their support and the resources for this research. This study was also funded by grant PI19/00878 awarded to M. T within the State Plan for R+D+I 2013-2016 (National Plan for Scientific Research, Technological Development and Innovation 2008-2011) and co-financed by the ISCIII-Deputy General Directorate for Evaluation and Promotion of Research - European Regional Development Fund “A way of Making Europe” and Instituto de Salud Carlos III FEDER. M. T. was financially supported by the Miguel Servet Research Programme (SERGAS and ISCIII). I. B. was financially supported by pFIS program (ISCIII, FI20/00302). O. P. was financially supported by a grant IN606A-2020/035 (GAIN, Xunta de Galicia). The author acknowledge CESGA (www.cesga.es) in Santiago de Compostela, Spain, for providing access to computing facilities.

The authors declare that there are no conflicts of interest.

